# The effect of NMDA-R antagonist, MK-801, on neuronal mismatch along the auditory thalamocortical pathway

**DOI:** 10.1101/636068

**Authors:** Gloria G Parras, Catalina Valdés-Baizabal, Lauren Harms, Patricia Michie, Manuel S Malmierca

## Abstract

Efficient sensory processing requires that the brain is able to maximize its response to unexpected stimuli, while suppressing responsivity to expected events. Mismatch negativity (MMN) is an auditory event-related potential that occurs when a regular pattern is interrupted by an event that violates the expected properties of the pattern. MMN has been found to be reduced in individuals with schizophrenia in over 100 separate studies, an effect believed to be underpinned by glutamate N-methyl-D-aspartate receptor (NMDA-R) dysfunction, as it is observed that NMDA-R antagonists also reduce MMN in healthy volunteers. The aim of the current study is to examine this effect in rodents. Using single unit recording in specific auditory areas using methods not readily utilized in humans, we have previously demonstrated that neuronal indices of rodent mismatch responses recorded from thalamic and cortical areas of the brain can be decomposed into a relatively simple repetition suppression and a more sophisticated prediction error process. In the current study, we aimed to test how the NMDA-R antagonist, MK-801, affected both of these processes along the rat auditory thalamocortical pathway. We found that MK-801 had the opposite effect than expected, and enhanced thalamic repetition suppression and cortical prediction error. These single unit data correlate with the recordings of local field responses. Together with previous data, this study suggests that our understanding of the contribution of NMDA-R system to MMN generation is far from complete, and also has potential implications for future research in schizophrenia.

**Significance Statement:** In this study, we demonstrate that an NMDA-R antagonist, MK-801, differentially affects single neuron responses to auditory stimuli along the thalamocortical axis by increasing the response magnitude of unexpected events in the auditory cortex and intensifying the adaptation of responses to expected events in the thalamus. Thus, we provide evidence that NMDA-R antagonists alter the balance between prediction-error and repetition suppression processes that underlie the generation of mismatch responses in the brain, and these effects are differentially expressed at different levels of auditory processing. As effects of MK-801 were in the opposite direction to our expectations, it demonstrates that our understanding of role of NMDA-R in synaptic plasticity and the neural processes underpinning MMN generation are far from complete.

## INTRODUCTION

Mismatch negativity (MMN) is an auditory event-related potential (ERP) that occurs when an unexpected stimulus (deviant, DEV) interrupts a train of regular stimuli (standards, STD) in an *oddball* sequence. MMN is commonly quantified as the difference between the size of the DEV and STD ERPs (Näätänen *et al.*, 1978).

The predictive coding framework has emerged as an appealing model of MMN (Randeniya *et al.*, 2018) and of how sensory information is processed. According to this model, the brain constantly generates top-down predictions from any regular ascending input that is compared with sensory bottom-up signals. Neural responses to stimuli that match predictions are suppressed, whereas unexpected stimuli discrepant with the prediction generate an enhanced error signal (Friston, 2005; Michie *et al.*, 2016; Carbajal & Malmierca, 2018). NMDA-R dependent plasticity is believed to underpin the capacity of the brain to adjust internal predictions and use memory of recent past inputs to anticipate future stimuli (Wacongne, 2016).

There are two mechanisms underlying the MMN difference signal according to the predictive coding model. First, MMN could reflect *repetition suppression*. When the same stimulus is repeatedly presented, neuronal populations sensitive to that stimulus undergo adaptation and neural responses decrease (Bendixen *et al.*, 2007). MMN could also reflect a process of *prediction error*, where the sensory memory of regular stimuli establishes a predictive model. Violation of this prediction upon presentation of an unexpected DEV stimuli results an enhanced neural response. Prediction error has been observed in human and rodent surface recordings when suitable control conditions have been included in the design of sound sequences (Nakamura *et al.*, 2011; Harms *et al.*, 2014; Parras *et al.*, 2017; Kurkela *et al.*, 2018). Essentially, there is evidence in both humans and rodents that MMN receives contributions from both prediction error and repetition suppression at various levels of the auditory system (Parras *et al.*, 2017; Ishishita *et al.*, 2018).

MMN reduction occurs in healthy volunteers administered an NMDA-R antagonist (Umbricht *et al.*, 2000) and has been reported in over 100 separate reports in individuals diagnosed with schizophrenia (Todd et al., 2013), leading to assertions that MMN indexes the functional state of NMDA-R neurotransmission (Umbricht *et al.*, 2002). The schizophrenia findings fit with current views that NMDA-R hypofunction contributes to the neuropathology of the disorder (Krystal *et al.*, 2005; Javitt et al., 1996). Mismatch like responses (MMRs) in animal models have shown a similar sensitivity to NMDA-R antagonism (Javitt *et al.*, 1996; Siegel *et al.*, 2013; Harms, 2016; Featherstone *et al.*, 2018).

Our primary interest here is whether NMDA-R antagonists differentially affect repetition suppression and prediction error at the single unit level at both the thalamic level and cortex. While previous studies have demonstrated that MMN-like responses in rodents are altered by NMDA-R antagonists (Ehrlichman *et al.*, 2008; Tikhonravov *et al.*, 2008, 2010; Sivarao *et al.*, 2014), only one report has examined the impact on prediction error component of the MMN in surface recordings (Harms *et al.*, 2018). There are no published data on the effects of NMDA-R antagonists on repetition suppression. More importantly, there are no reports that have examined the effects of NMDA-R antagonists on single-unit activity and local field potential recordings from the thalamus and auditory cortex.

Thus, it is unknown (i) whether there are differential effects of NMDA-R antagonism on prediction error as opposed to repetition suppression at the single unit or local field potential level, and (ii) the regional specificity of where effects of NMDA-R antagonists occur: in the lemniscal *vs*. non-lemniscal auditory areas, or the thalamus *vs*. cortex. Therefore, we used an acute exposure to a low dose of MK-801 to examine the impact of NMDA-R antagonism on individual responses of lemniscal and non-lemniscal thalamocortical neurons while auditory oddball, many standards and cascade control sequences were presented (figure 1a-b). This design allowed us to delineate effects on repetition suppression *vs*. prediction error (figure 1c)(Opitz *et al.*, 2005; Ruhnau *et al.*, 2012; Harms, 2016; Parras *et al.*, 2017). Unexpectedly, we found that MK-801 increased repetition suppression in the thalamic regions, enhanced prediction error in the cortex and increased neuronal mismatch at both locations.

**Figure 1.**
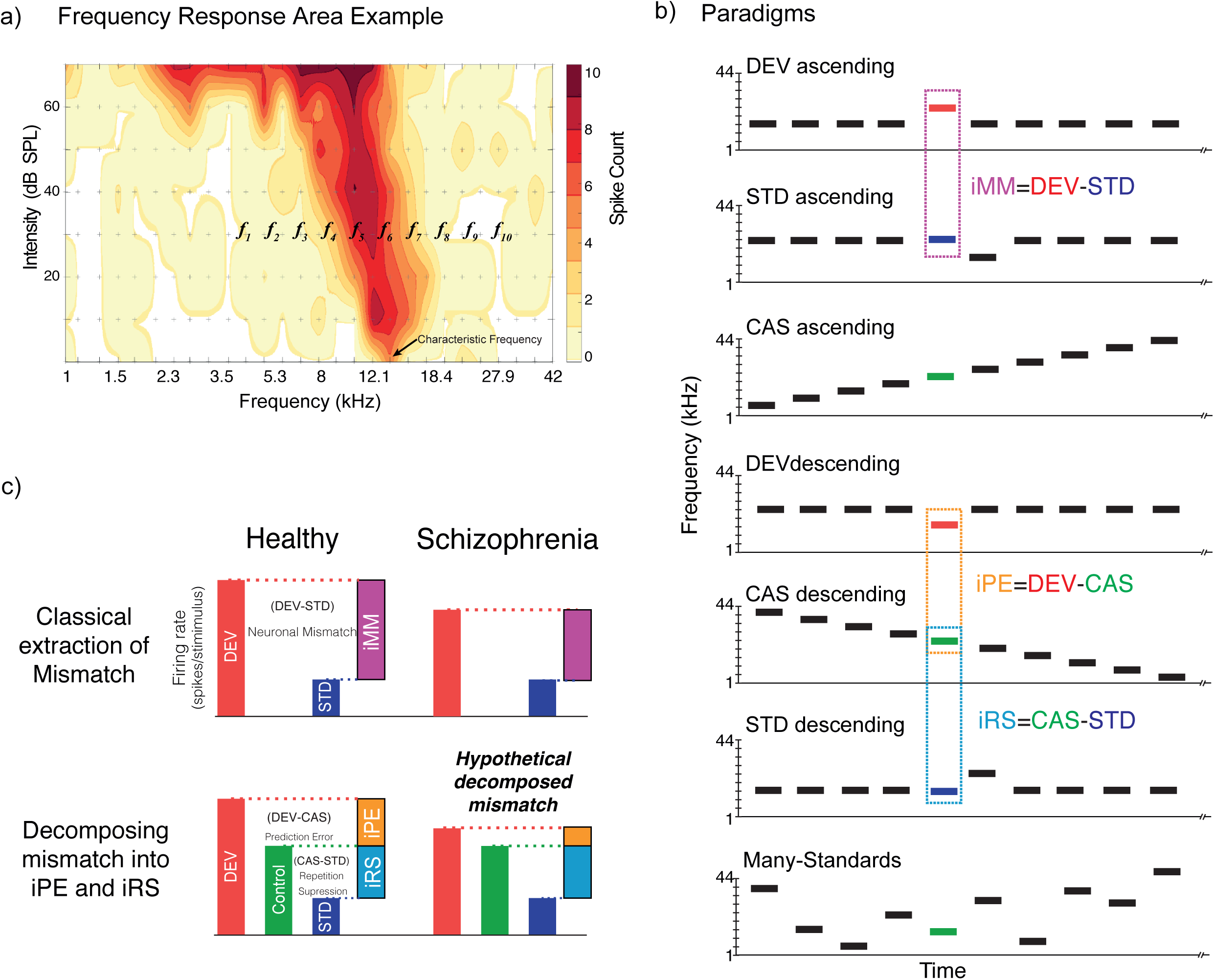
Experimental design. **a)** Frequency response area example, with a representation of the ten selected tones to build the experimental paradigms. **b)** Stimulation sequences, the same tone could be presented in different experimental paradigms, thus we can compare same tone in different contexts to control adaptation and deviance detection; and conform the indices of neuronal mismatch (iMM), prediction error (iPE) and repetition suppression (iRS). Note that ascending and descending tones will be compared to the control ascending or descending, respectively. **c)** Sketch of summary results of mismatch responses for healthy and schizophrenia subjects under the classical analysis of mismatch. Second row decomposition of neuronal mismatch, under the assumption of predictive coding framework in healthy subjects, and the hypothetical decomposition of neuronal mismatch into prediction error and repetition suppression in schizophrenia.

## MATERIAL AND METHODS

Experiments were performed on 48 (control=25; MK-801=23) adult, female Long-Evans rats with body weights between 200-250g (aged 9 to 15 weeks). All experimental procedures were performed at the University of Salamanca, and all procedures and experimental protocols were in accordance with the guidelines of the European Communities Directive (86/609/EEC, 2003/65/EC and 2010/63/EU) and the RD 53/2013 Spanish legislation for the use and care of animals. All the details of the study were approved by the Bioethics Committee of the University of Salamanca (ref# USAL-ID-195).

### Surgical procedures

Anesthesia was induced and maintained with urethane (1.5g/kg, i.p), with supplementary doses (0.5g/kg, i.p) given as needed. Dexamethasone (0.25mg/kg) and atropine (0.1mg/kg) were administered at the beginning of the surgery to reduce brain edema and bronchial secretions, respectively. Isotonic glucosaline solution was administered periodically (5-10ml every 6-8h, s.c) to avoid dehydration. During all experimental procedures, animals were artificially ventilated, and CO_2_ and temperature monitored (Pérez-González *et al.*, 2012; Duque & Malmierca, 2015).

The initial procedure was the same in each case, and the subsequent procedures differed only in the craniotomy location, and the placement/orientation for the recording electrode (animals per group/location: control MGB=16, AC=9; MK-801 MGB=15, AC=8). For MGB recordings, a craniotomy (∼2×2mm, from −5 to −6.5mm bregma and - 3.5mm lateral) was performed in the left parietal bone, dura was removed, and the electrode advanced in a vertical direction (Antunes *et al.*, 2010). For AC recordings, the skin and muscle over the left temporal bone was retracted and a 6×5mm craniotomy was performed (between −2 and −6 from Bregma) over the temporal bone (Nieto-Diego & Malmierca, 2016) dura was removed and the area was covered with a thin, transparent layer of agar to prevent desiccation and stabilize recordings. Electrodes for AC recording were inserted using a triple axis micromanipulator (Sensapex), forming a 30° angle with the horizontal plane, to penetrate through all cortical layers of the same cortical column.

For this study, animals in MK-801-treated group receive a systemic intraperitoneal injection (0.1mg/kg) of a noncompetitive NMDA-R antagonist (MK-801 hydrogen maleate, M107 Sigma-Aldrich). Control animals did not receive any injection.

### Electrophysiological recording procedures

During all procedures, animals were placed in a stereotaxic frame fixed with hollow specula ear bars that housed the sound delivery system. One single neuron and local field potential (LFP) was recorded at a time, using the same tungsten electrode (1-4MΩ) inserted into a single auditory station (MGB or AC) in each individual animal. The signal recorded was pre-amplified (1000x) and band-pass filtered (1-3kHz) with a medusa preamplifier (TDT). This analog signal was digitalized 12k sampling rate and further band-pass filtered (TDT-RX6) separately for spikes (500Hz-3kHz) and LFP (3-50Hz). We used short trains of white noise bursts (30 ms, 5 ms rise-fall ramps) to search for neuronal activity. To prevent neuronal adaptation during the search, some parameters (frequency and intensity) and stimulus type (white noise, pure tone) were manually varied. Once a single neuron was isolated a frequency-response area (FRA) of the response magnitude for each frequency/intensity combination was first computed (Figure 1a). A randomized sequence of pure tones (from 1 to 44 KHz) was presented at a rate of 4Hz, with varying frequency and intensity, and with 3 repetitions of all tones.

For each animal treated with MK-801 the first single neuron was recorded ∼15 min after the drug injection (Vezzani *et al.*, 1989). Ten evenly-spaced pure tones (0.5 octaves separation) at a fixed sound intensity (usually 20-30dB above the threshold) were selected to each neuron recorded to create the control sequences, cascades and many-standard (Ruhnau *et al.*, 2012; Parras *et al.*, 2017), and additionally, adjacent pairs of them were used to present various oddball sequences (Figure 1b). All sequences were 400 tones in length (75ms duration, 5ms rise-fall ramp and 250ms interstimulus interval), each tone in the control sequences was played 40 times, with the same overall presentation rate as deviants in the oddball sequence.

Oddball sequences were used to test the specific contribution of deviant tones in an adaptation context. An oddball sequence consisted of a repetitive tone (standard 90% probability), occasionally replaced by a tone of a different frequency (deviant 10% probability), in a pseudorandom manner. We used two types of control sequences: the many-standard and cascade sequences. Both containing the same 10 frequencies but differing in the order of presentation. The many-standard control was randomly presented, mimicking the presentation rate and the unpredictability of the deviant tones. While cascades were played always in the same presentation order, ascending or descending in frequency. Hence the cascade contains a regularity, mimic the presentation rate of deviant sounds but in a predictable context and consequently do not violate a regularity. These four conditions, and by extension responses to them, will be denoted as deviant (DEV), standard (STD), cascade (CAS) and many-standard. Finally, if the neuron could be held for long enough, the same protocol was repeated for different frequencies and/or intensity.

### Anatomical location

For MGB recording localization, at the end of each tract and experiment, two electrolytic lesions were made to mark the end and the beginning of the auditory signal (figure 6a). Then, animals were given a lethal dose of sodium pentobarbital and perfused transcardially with phosphate buffered saline (0.5% NaNO_3_ in Phosphate Buffered Saline) followed by a fixative mix of 1% paraformaldehyde and 1% glutaraldehyde). After fixation and dissection, the brain was cryoprotected in 30% sucrose and sectioned into 40μm slices. Sections were Nissl stained with 0.1% cresyl violet. Recording sites were marked on images from an adult rat brain atlas (Paxinos & Watson, 2013) and neurons that were recorded from were assigned to one of the main divisions of the MGB (dorsal, medial or ventral). This information was complemented and confirmed by the stereotaxic coordinates as well as the depth of the neuron within a tract.

For the AC experiments, a magnified picture (25x) of the exposed cortex and the Bregma references was taken at the end of the surgery with a digital single lens reflex camera (D5100, Nikon) coupled to the surgical microscope (Zeiss). The picture was overlapped to guide and mark each electrode placement into a micrometric grid (250-500 ∼μm spacing; figure 6b). Then we performed several tracts recording multi-unit activity frequency response area (FRA), the characteristic frequency arise from each FRA was placed over the picture, resulting in a characteristic frequency map of each animal. Boundaries were identify following the changes in the tonotopic gradient: high-frequency reversal between the ventral and anterior auditory fields (rostrally), low-frequency reversal between primary and posterior auditory field (dorsocaudally) and high-frequency reversal between ventral and suprarhinal auditory field (ventrally) (Nieto-Diego & Malmierca, 2016). Then, each recording was located in one of these five fields. Nevertheless, the map was complemented during all electrophysiological recording session with the characteristic frequency of each new tract.

### Statistical analysis

All the data analyses were performed with MatlabTM software, using the built-in functions, the Statistics and Machine Learning toolbox, or custom scripts and functions developed in our laboratory. To test for significant excitatory responses to tones we used a Monte Carlo approach, simulating 1000 peri-stimulus time histogram (PSTH) using a Poison model with a constant firing rate equal to the spontaneous firing rate. A null distribution of baseline-corrected spike counts was generated from this collection of PSTH. Lastly, the *p* value of the baseline-corrected spike count was empirically computed as *p* = (g+ 1)/(N+ 1), where g is the count of null measures greater than or equal to baseline-corrected spike count, and N=1000 is the size of the null sample. Finally, we only included in the analysis neuron/frequency combinations with significant excitatory response (*p* > 0.05) after the baseline-corrected spike count to at least one of the conditions (DEV, STD, CAS). PSTH were used to estimate the spike-density function (SDF) over the time, showing action potential density over time (in action potentials per second) from −75 to 250ms around stimulus onset, for the 40 trials available for each tone and condition (DEV, STD, CAS), smoothed with a 6ms gaussian kernel (“ksdensity” function in Matlab) in 1ms steps. The baseline spontaneous firing rate was determined as the average firing rate during the 75ms preceding stimulus onset.

The excitatory response was measured as the area below the SDF and above the baseline spontaneous firing rate, between 0 and 180ms after stimulus onset (positive area patches only, to avoid negative response values). This measure will be referred to as “baseline-corrected spike count”.

Baseline-corrected spike count responses of a neuron to the same tone in the three conditions (DEV, STD, CAS) were normalized using the formulas:

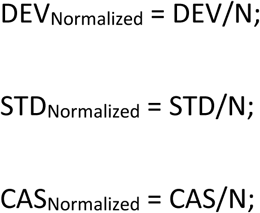

Where 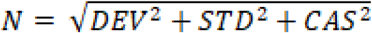, is the Euclidean norm of the vector (DEV, STD, CAS) defined by the three responses. Normalized values were the coordinates of a 3D unit vector (DEV_Normalized_, STD_Normalized_, CAS_Normalized_) with the same direction of the original vector (DEV, STD, CAS), and thus the same proportions between the three response measures. This normalization procedure always results in a value ranging 0–1, and has a straightforward geometrical interpretation.

From these normalized responses, indices of neuronal mismatch (iMM), repetition suppression (iRS), and prediction error (iPE) were computed as:

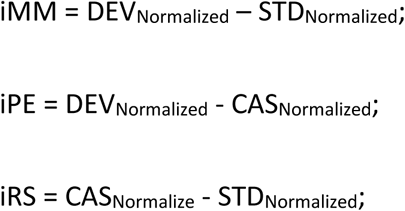

These indices, consequently, always range between −1 and 1, and provide the following quantitative decomposition of neuronal mismatch into repetition suppression and prediction error: iMM = iRS + iPE. To test these indices over time, we divided the whole response into 12 time windows, 20ms width, from −50 to 190ms with respect to the stimulus onset. Then, we compared each time window against zero using a sign-rank test, false discovery rate (FDR=0.1) corrected for the 12 windows.

For the analysis of the LFP signal, we aligned the recorded wave to the onset of the stimulus for every trial, and computed the mean LFP for every recording site and stimulus condition (DEV-LFP, STD-LFP and CTR-LFP), as well as the differences between them, resulting in the three LFP-indices: “neuronal mismatch” (MM-LFP = DEV-LFP – STD-LFP), “prediction error” (PE-LFP = DEV-LFP – CAS-LFP) and “repetition suppression” (RS-LFP = CAS-LFP – STD-LFP). Then, grand-averages were computed for all conditions and auditory station separately. The *p* value of the grand-averaged for the three LFP-indices (MM-LFP, PE-LFP and RS-LFP) was determined for every time point with a two-tailed *t* test (FDR corrected).

Our data set was not normally distributed, so we used distribution-free (non-parametric) tests. These included the Wilcoxon signed-rank test and Friedman test (for baseline-corrected spike counts, normalized responses, indices of neuronal mismatch, repetition suppression and prediction error). Only the difference wave for the LFPs was tested using a *t*-test, since each LFP trace is itself an average of 40 waves. For multiple comparison tests, *p* values were FDR corrected using the Benjamini-Hochberg method. Linear models were used to test for significant average iMM, iPE and iRS within each auditory station. Significant effects of station, pathway, and interactions between them were fitted using the ‘fitlm’ function in Matlab, with robust options. To estimate final sample sizes required for the observed effects after the initial exploratory experiments, we used the ‘sampsizepwr’ function in Matlab adjusted for the iPE for each region, to obtain a statistical power of 0.8 for this index. Sample sizes were enlarged with additional experiments until they were just greater than the minimum required (number of points recorded, and the minimum required for each station; see Table 1).

**Table 1.**
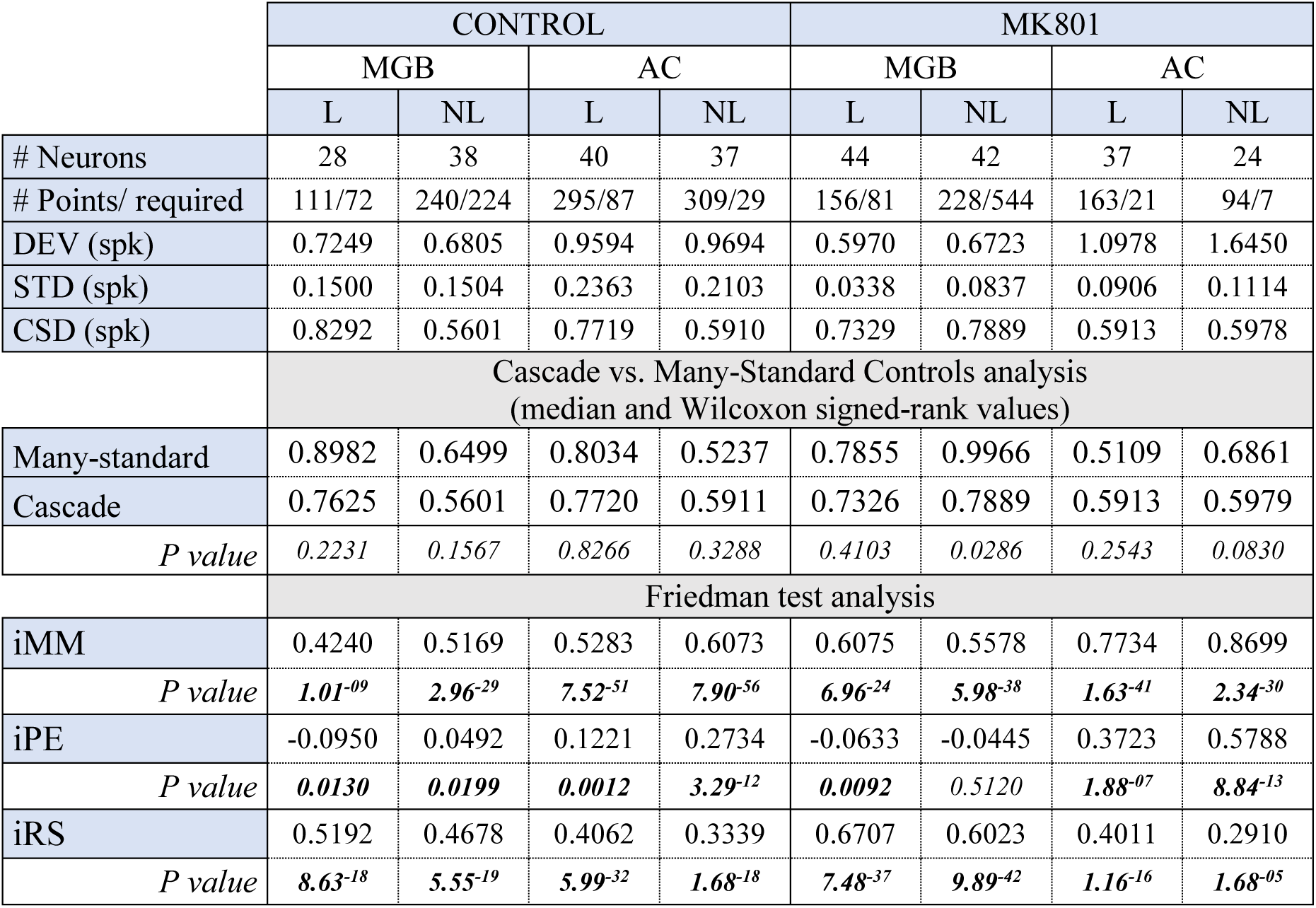
**Spike population analysis for** each experimental group and auditory station independently: First row, number of recorded neurons; second row number of tested neuron/frequency combinations (points), along with estimated minimum sample size (of points) required for a statistical power (See Methods). Followed by median values for base-line corrected spike count (spikes) to the different conditions. Comparative analysis for control paradigms, median values and Wilcoxon signed-rank test values for each station and group. Median indices of neuronal mismatch (iMM), prediction error (iPE) and repetition suppression (iRS), and their corresponding *p* value.

|  | CONTROL |  |  |  | MK801 |  |  |  |
| --- | --- | --- | --- | --- | --- | --- | --- | --- |
|  | MGB |  | AC |  | MGB |  | AC |  |
|  | L | NL | L | NL | L | NL | L | NL |
| # Neurons | 28 | 38 | 40 | 37 | 44 | 42 | 37 | 24 |
| # Points/ required | 111/72 | 240/224 | 295/87 | 309/29 | 156/81 | 228/544 | 163/21 | 94/7 |
| DEV (spk) | 0.7249 | 0.6805 | 0.9594 | 0.9694 | 0.5970 | 0.6723 | 1.0978 | 1.6450 |
| STD (spk) | 0.1500 | 0.1504 | 0.2363 | 0.2103 | 0.0338 | 0.0837 | 0.0906 | 0.1114 |
| CSD (spk) | 0.8292 | 0.5601 | 0.7719 | 0.5910 | 0.7329 | 0.7889 | 0.5913 | 0.5978 |
|  | Cascade vs. Many-Standard Controls analysis<br>(median and Wilcoxon signed-rank values) |  |  |  |  |  |  |  |
| Many-standard | 0.8982 | 0.6499 | 0.8034 | 0.5237 | 0.7855 | 0.9966 | 0.5109 | 0.6861 |
| Cascade | 0.7625 | 0.5601 | 0.7720 | 0.5911 | 0.7326 | 0.7889 | 0.5913 | 0.5979 |
| *P value* | 0.2231 | 0.1567 | 0.8266 | 0.3288 | 0.4103 | 0.0286 | 0.2543 | 0.0830 |
|  | Friedman test analysis |  |  |  |  |  |  |  |
| iMM | 0.4240 | 0.5169 | 0.5283 | 0.6073 | 0.6075 | 0.5578 | 0.7734 | 0.8699 |
| *P value* | **1.01<sup>-09</sup>** | **2.96<sup>-29</sup>** | **7.52<sup>-51</sup>** | **7.90<sup>-56</sup>** | **6.96<sup>-24</sup>** | **5.98<sup>-38</sup>** | **1.63<sup>-41</sup>** | **2.34<sup>-30</sup>** |
| iPE | -0.0950 | 0.0492 | 0.1221 | 0.2734 | -0.0633 | -0.0445 | 0.3723 | 0.5788 |
| *P value* | **0.0130** | **0.0199** | **0.0012** | **3.29<sup>-12</sup>** | **0.0092** | **0.5120** | **1.88<sup>-07</sup>** | **8.84<sup>-13</sup>** |
| iRS | 0.5192 | 0.4678 | 0.4062 | 0.3339 | 0.6707 | 0.6023 | 0.4011 | 0.2910 |
| *P value* | **8.63<sup>-18</sup>** | **5.55<sup>-19</sup>** | **5.99<sup>-32</sup>** | **1.68<sup>-18</sup>** | **7.48<sup>-37</sup>** | **9.89<sup>-42</sup>** | **1.16<sup>-16</sup>** | **1.68<sup>-05</sup>** |

To analyze the time course of adaptation we computed an averaged time course for all the standard stimuli presented. Then, we fitted a power law function with a three parameters model, *y(t)=a·t^b^+c*, where *a* indicates the responses beginning or the first spike strength; *b* the sensitivity to repetitive stimuli, or the adaptation velocity, and *c* the steady-state response. R^2^ values indicated that the model fits very well for standard responses in both groups, explaining between 60% and 78% of the response variability within all regions.

To analyze spikes differences between MK-801 and control group we computed the median values for each condition tested (DEV, STD and CAS) and their differences (iMM, iRS and iPE) and calculated a ranksum test. To compare each time window between groups a two-sample *t*-test (from 0 to 200ms, Bonferroni corrected for 200 comparisons with family-wise error rate FWER< 0.05) was used for the SDF and LFPs to each stimulus condition and indices, using the ‘ttest2’ function in Matlab, for every time point.

## RESULTS

We recorded a total of 290 well isolated neurons, 143 from the control group and 147 from the MK-801-treated group. One single neuron and local field potential (LFP) was simultaneously recorded at a time, using the same tungsten electrode. Recordings were performed in the medial geniculate body (MGB) and in the auditory cortex (AC) while playing oddball and control sequences (many-standards and cascade: CAS) in anesthetized rats. Since we found no statistically significant differences between the use of the cascade and many-standards sequences for the control group and MK-801 group, except for the MGB_NL_ from the MK-801 group (table 1), the CAS sequence was chosen to control for repetition effects. This is because the CAS paradigm not only controlled for the presentation rate of the deviant stimuli, but also the frequency difference (ascending or descending) between standards and deviants in the oddball sequences.

### Effects of MK-801 on the neuronal firing rate

MK-801 injection significantly reduced the responses to STD tones within all regions. By contrast, for responses to the DEV tones, we observed a significant increment in responses in AC but not for the MGB. When the firing rate of the cascade sequence was considered, MK-801 differentially affected the AC and MGB such that CAS responses were significantly increased in the MGB_NL_ but decreased in AC. These results reveal a differential effect of MK-801 on the refractoriness and salience of infrequent events at the single neuron level (Figure 2a, table 2).

**Figure 2.**
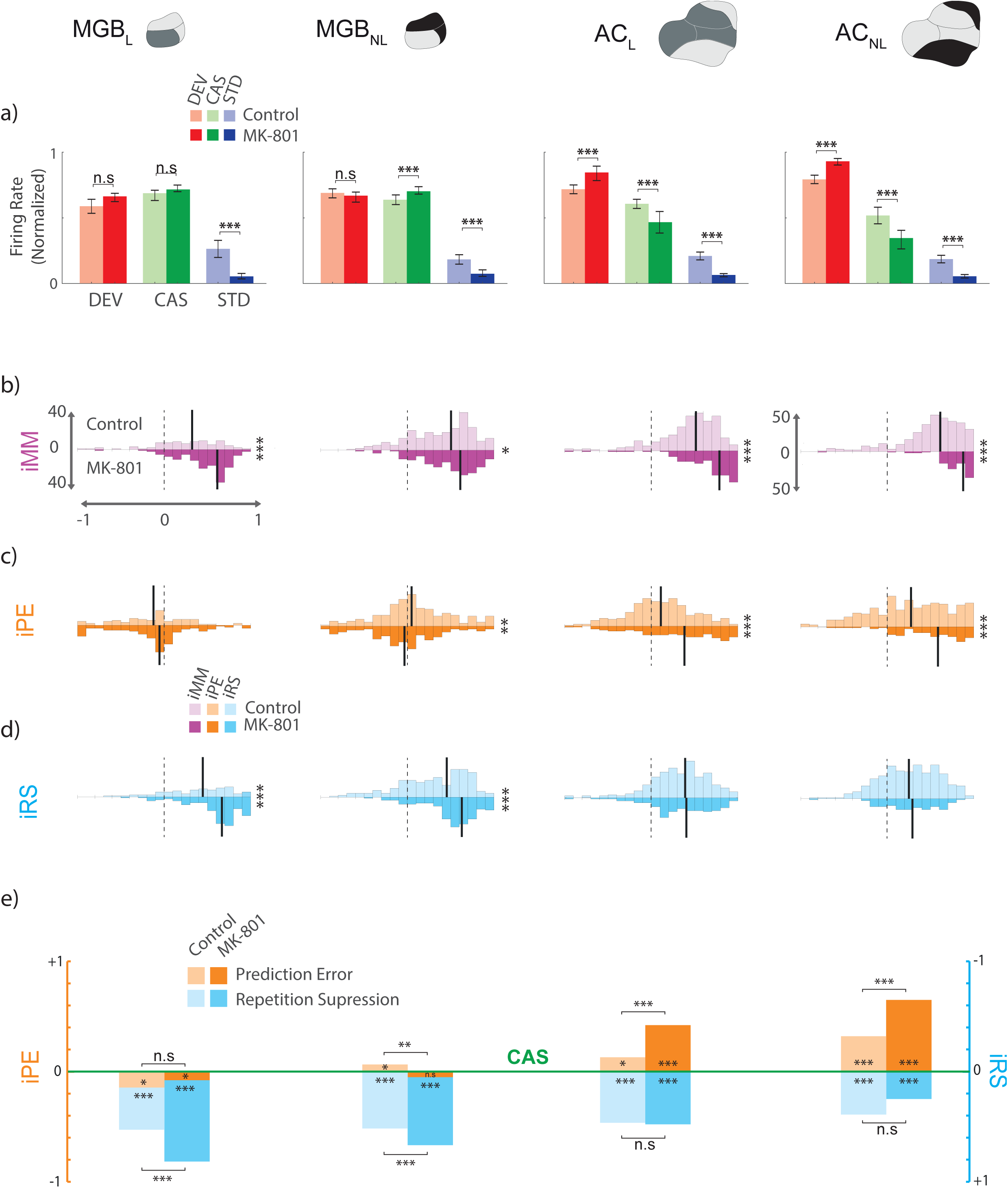
Single neuron spikes population analysis. Results for firing rate analysis and their computed differences along the thalamocortical axis. **a)** Boxplot of median normalized responses for deviants (red), cascade (green) and standard (blue) for each group, control (light colors) and MK801 (bright colors), within each station and the statistical significance between groups (Wilcoxon signed-rank test, * *p*<0.05, \*\**p*<0.001, \*\*\**p*<0.000). **b-d)** Indices histograms displayed in a mirror-like manner for the two groups (controls upper and in light colors; MK801 under and in bright colors), showing the distribution of the three indexes for each neuronal response (ranging between −1 and +1, dotted lines indicate index=0). Vertical solid lines indicate their medians and the significant difference between groups is noted at the right of each histogram block. **e)** Median indices of Prediction Error (orange) and **d)** Repetition Suppression (blue), represented with respect to the baseline set by the cascade control (green line). Thereby, iPE upwards-positive while iRS is downwards-positive. Each median index corresponds to differences between normalized responses in a). Asterisks inside bars denote statistically significance of these indices against zero (Friedman test), while asterisks outside bars denote statistically significance between groups (Wilcoxon signed-rank test, * *p*<0.05, \*\**p*<0.001, \*\*\**p*<0.000).

**Table 2:** Firing Rate Comparisons. **Comparative analysis between control and MK-801 group.** Median spikes to the former measures responses to standard (STD), deviant (DEV) and cascade (CAS) tones, and their corresponding *p* value (ranksum test). Similarly, medians and their associates *p* value for the index of mismatch (iMM), index of prediction error (iPE) and index of repetition suppression (iRS).

|  | MGB |  | AC |  |
| --- | --- | --- | --- | --- |
|  | L | NL | L | NL |
| STD_control | 0.1912 | 0.1717 | 0.1990 | 0.1856 |
| STD_MK-801 | 0.0537 | 0.1047 | 0.0689 | 0.0547 |
| *P value* | **<0.000** | **<0.000** | **<0.000** | **<0.000** |
| DEV_control | 0.6153 | 0.6887 | 0.7274 | 0.7930 |
| DEV_MK-801 | 0.6611 | 0.6620 | 0.8424 | 0.9247 |
| *P value* | 0.2309 | 0.2641 | **<0.000** | **<0.000** |
| CAS_control | 0.7104 | 0.6395 | 0.6052 | 0.5196 |
| CAS_MK-801 | 0.7244 | 0.7063 | 0.4701 | 0.3458 |
| *P value* | 0.1169 | **<0.000** | **<0.000** | **<0.000** |
| iMM_control | 0.3675 | 0.4897 | 0.5124 | 0.5781 |
| iMM_MK-801 | 0.5197 | 0.5221 | 0.7269 | 0.8088 |
| *P value* | **<0.000** | **0.0409** | **<0.000** | **<0.000** |
| iPE_control | -0.0670 | 0.0513 | 0.1060 | 0.2697 |
| iPE_MK-801 | -0.0502 | -0.0444 | 0.3592 | 0.5652 |
| *P value* | 0.9622 | **0.0035** | **0.000** | **<0.000** |
| iRS_control | 0.5300 | 0.3672 | 0.3350 | 0.2935 |
| iRSdrug | 0.6812 | 0.5632 | 0.3514 | 0.2765 |
| *P value* | **<0.000** | **<0.000** | 0.5800 | 0.8934 |

### Effects of MK-801 on neuronal mismatch and its components

Next we analyzed the differences between these normalized responses and computed three indexes (ranging between −1 and +1): 1) the index of neuronal mismatch (iMM=DEV-STD), similar to the typical SSA index used in previous single neurons studies; 2) the index of prediction error (iPE= DEV-CAS), that shows the relative enhancement of DEV tones compared with CAS tones and 3) the index of repetition suppression (iRS=CAS-STD) that reflects the level of response suppression due to the repetition effect, and is obtained by comparing the normalized responses to CAS and STD. It should be noted that the iMM is the sum of iRS and iPE (iMM=iRS+iPE).

The analysis of the iMM after the injection of MK-801 demonstrated that iMM values are significantly different from zero for all recording sites (figure 2b, table 1: Friedman test). But when comparisons between groups were considered, the analysis revealed that MK-801 increased the neuronal iMM (figure 2b-iMM; table 2). As described above, these changes are largely due to a reduced response to STD tones in all recording locations and an enhanced response to DEV in the AC.

Since iMM=iRS+iPE, an important advantage of these metrics is that we can determine how much of the mismatch index is due to the regularity of the context (RS) and/or to the occurrence of an infrequent event (PE). Thus, to determine which of these two components of the iMM is affected by MK-801, we computed the indices of iPE and iRS separately.

Interestingly, MGB neurons in the MK-801 group did not show any sign of genuine deviance detection, as iPE values were almost zero and negative. While both AC showed a significant positive iPE (figure 2c; iPE values in table 1). When comparison between groups were analyzed an increased iPE for the MK-801 group in the AC were found, and even a further decreased iPE for the MGB_NL_ in the MK-801 group (figure 2c and e light and bright oranges; iPE in table 2). These data suggest that the MK-801 produces an augmentation of saliency for novel stimuli processed in the AC.

Yet, the detection of rare or novel stimuli requires the establishment of a regular context or pattern. Therefore, we were also interested to find out if the refractoriness due to regularity was altered by MK-801. We calculated the iRS by assessing the response of the same tone when it was presented as CAS, with a 10% probability in a regular pattern and presented as STD with a probability of 90%, within an oddball paradigm, so it is in a much more regular context (Harms et al., 2014; Ruhnau et al., 2012). In both cases, we assume some level of regularity adaptation, but only a genuine repetition suppression can be determined if the responses to STD tones are lower than responses to CAS. Our results demonstrate that there is a significant repetition suppression effect in the MK-801 group along the thalamocortical pathway (figure 2d bright blue; iRS in table 1). The analysis also revealed that MK-801 produced a significant increase in repetition suppression at thalamic level but did not affect repetition suppression in the AC when compared with controls (figure 2d-e light and bright blues; results in table 2).

These results show that the auditory thalamus and cortex differ in the way repetition effects and prediction errors are processed. To confirm this hypothesis and considering that we have previously found an increase in the level of iPE along the thalamocortical hierarchy in awake and anesthetized animals (Parras et al., 2017), we fitted a linear model to determine the effect of MK-801 on the degree of increase in iPE along the auditory hierarchy relative to the control (urethane) condition. Using station (S: MGB vs. AC), drug (D: Control vs. MK-801) and pathway (P: Lemniscal *vs.* Non-lemniscal) and their interactions as factors, with control MGB_L_ as reference level for these factors, the fitted model is as follows:

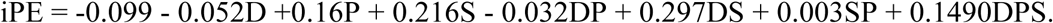

Next, we applied an ANOVA to this model and found a significant effect of station (F=209.24, *p*=1.21×10^-44^), drug (F=10.82, *p*=0.001), pathway (F=42.93, *p*=7.66×10^-11^) and for the interaction drug *x* station (F=37.9, *p*=9.41×10^-10^) but not for the interactions of drug *x* pathway (F=0.45, *p*=0.5011) nor station *x* pathway (F=0.004, *p*=0.9488), nor the three way interaction (F=2.37, *p*=0.1236). That is, there is a significant station effect (iPE is larger overall at AC than MGB), pathway (iPE is larger overall in the non-lemniscal than the lemniscal pathway), and drug (iPE is increased overall with MK801 vs control) and a drug x station interaction indicating that while iPE at AC is increased overall relative to MGB, the degree of iPE increase is moderated by MK801. Inspection of Figure 2e indicates that the significant station difference is in the direction of being larger under MK801 than urethane. These results indicate that indeed, the sensitivity to detect novel stimuli increases significantly more along the thalamocortical axis in the MK-801 than the control group (figure 2e, iPE in orange).

We also fitted a similar linear model to iRS. The resulting model was:

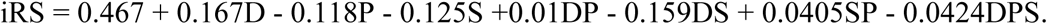

ANOVA demonstrated a significant effect for drug (F=26.42, *p*=3.08×10^-7^), station (F=80.52, *p*=7.89×10^-19^), pathway (F=24.65, *p*=7.6×10^-7^) and the interaction of drug *x* station (F=17.61, *p*=2.84×10^-5^), but not for the interactions drug *x* pathway (F=0.078, *p*=0.7790), station *x* pathway (F=1.45, *p*=0.2281) and the three way interaction (F=0.31, *p*=0.5759). Once again, these results indicate that while iRS was more marked at MGB than AC in general, this station difference was exacerbated with MK-801.

In summary, the changes described above demonstrate that NMDA-R antagonism has distinct effects on auditory scene analysis, as measured by the iPE and iRS, at different levels of the thalamocortical hierarchy.

### Effect of MK-801 on Spike-Density Function and indexes

Next, we sought to identify how MK-801 affected the temporal responses to auditory stimuli (DEV, STD and CAS) by comparing spike-density functions (SDF) to each condition between groups. Analysis revealed the latency of the main peak for the SDF to DEV tones was mostly unaffected by MK-801 in the MGB, but it was clearly delayed by 40 and 60 ms in the AC_L_ and AC_NL_, respectively. Furthermore, the magnitude of the SDF was altered at the AC and MGB_NL_, with the early component being reduced and the later sustained component being enhanced (figure 3a, horizontal white line for significant differences at *p*<0.05). When the STD tones were considered, we observed a distinct and significant decrease of the SDF mostly at the AC and only marginally at the subcortical levels (figure 3b). Finally, MK-801 affected mostly the initial responses to cascade tones at all regions, being reduced in the auditory cortex but was earlier and increased in MGB_L_ (figure 3c). The sustained portion of the SDF was only significantly increased in the MGB_NL_. Results show that MK-801 has a profound effect on the spike-density functions to DEV, STD and cascade stimuli.

**Figure 3.**
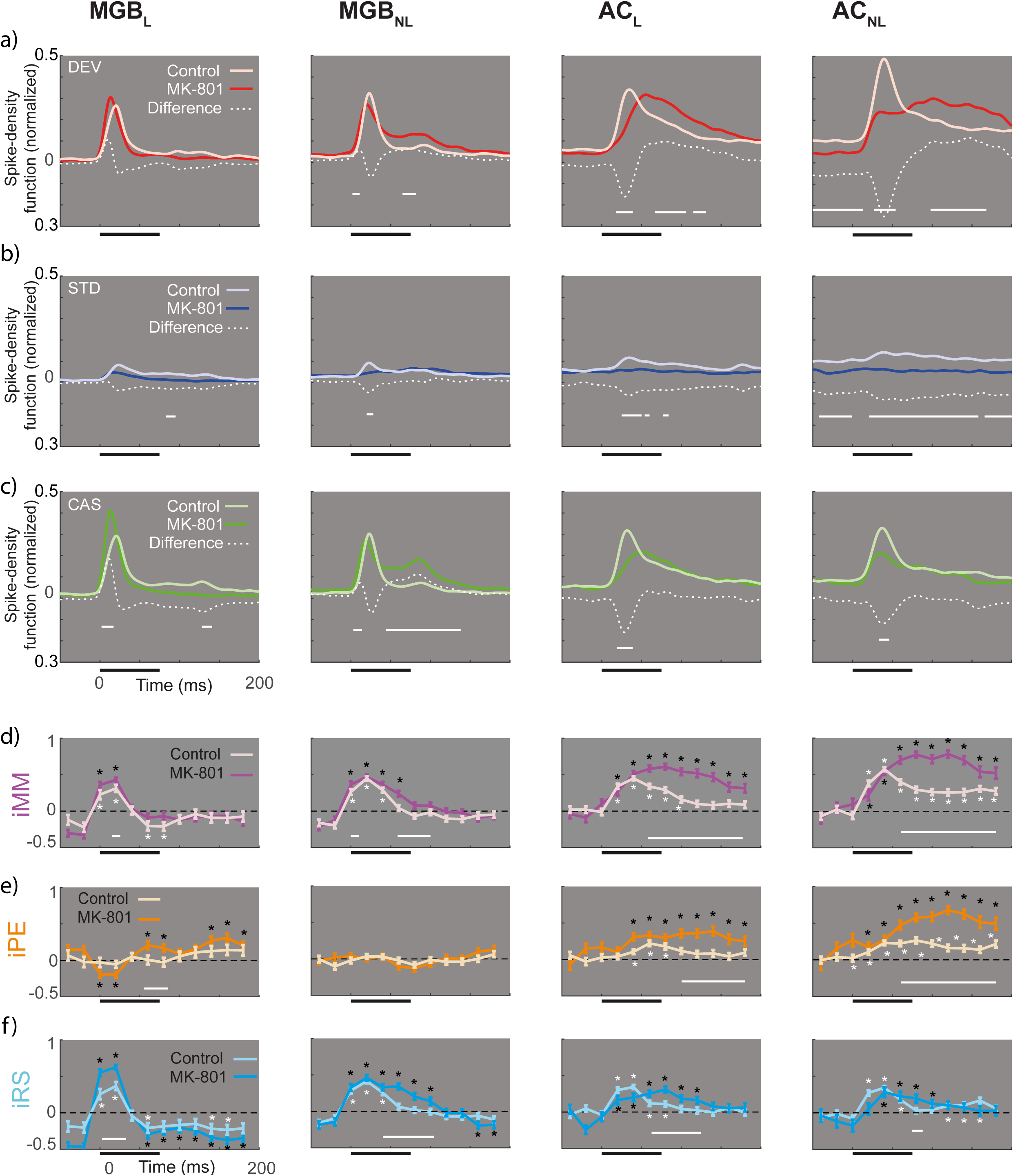
Spike Density Function. Peristimulus time histogram along the thalamocortical axis. **a-c)** Averaged firing rate profiles for each condition as normalized spike-density function (light colors for control and bright color for MK801 group), and their respective differences (white dotted lines). Solid horizontal white lines represent the time in which the difference between groups is significant (two-sample *t* test *p*<0.05, Bonferroni corrected). **d-f)** Indices over time computed for 12 intervals (from −50 to 190ms) compared against zero (signed-rank test and FDR corrected for 12 comparisons; * *p*<0.01) for each group (light colors for control and bright color for MK801 group). Solid white lines denote differences between groups across time intervalss (two-sample *t* test for each of the 12-time windows, *p*<0.05).

Next, we studied where and when the MK-801 effect on the neuronal indices of iMM, iPE and iRS was significantly different from control. Thus, we examined whether in each group independently (MK-810 and control) these indices are different from zero, *i.e.*, is there a significant iMM, iPE or iRS at each time point. Figure 3d-f highlights the significant time windows (*p*<0.01) with white and black asterisks for control and MK-801, respectively. The analysis revealed that under MK-801, there was a significant iMM along the thalamocortical axis (between 20-40ms for MGB_L_, 20-80ms in MGB_NL_ and from 20-190ms in both AC; Figure 3d, bright purple lines) and a significant iPE between 20 and 180ms in both AC, and a late iPE in the lemniscal thalamus between 60-80ms and 140-190ms (Figure 3e, bright orange lines). We also found significant thalamocortical iRS (figure 3f, bright cyan lines; between 20-40ms for MGB_L_, 0-100ms in MGB_NL_, from 20-120ms in AC_L_ and between 40-100ms in AC_NL_).

When we compared the two groups, the analysis revealed that MK-801 produced a significant enhancement of iMM and iPE at both AC subdivisions (*p*<0.000 for iMM between 60-190ms in both AC; and *p*<0.05 for iPE ranging between 100 and 190ms in AC_L_ and between 60-190ms in AC_NL_; white horizontal lines in Figures 3 d-f). By contrast, iRS was affected more in the MGB (*p*<0.000 between 5-35ms in MGB_L_; *p*<0.000 between 40-110ms in MGB_NL_; *p*<0.05 between 60-130ms in AC_L_; and *p*<0.05 at 80ms in AC_NL_; white horizontal lines in Figure 3f). Thus, MK-801 produces an increase of iMM and iPE mostly in the late time window in AC, while iRS is much affected in the MGB.

### MK-801 affects the dynamics of adaptation

Since MK-801 lowered and flattened responses to STD tones across the response window, we sought to assess the dynamics and the time course of adaptation (figure 4a). Results show that the control group (light gray arrows) exhibit a hierarchical timing for adaptation responses, becoming faster in higher order areas (from top to down, responses reach the half of the initial values at the fourth, ninth, twelfth and fourteenth standard tone, respectively). By contrast, results from the MK-801 group exhibited much faster adaptation dynamics (figure 4b; 50% of the initial response occurred at the third and second standard tones in MGB and AC, respectively; *b* values for control group: MGB_L_=-0.1769, MGB_NL_=-0.4174, AC_L_=-0.6824 and AC_NL_=-1.175; and for MK-801 group: MGB_L_=-0.8499, MGB_NL_=-0.8853, AC_L_=-1.712 and AC_NL_=-1.418).

**Figure 4.**
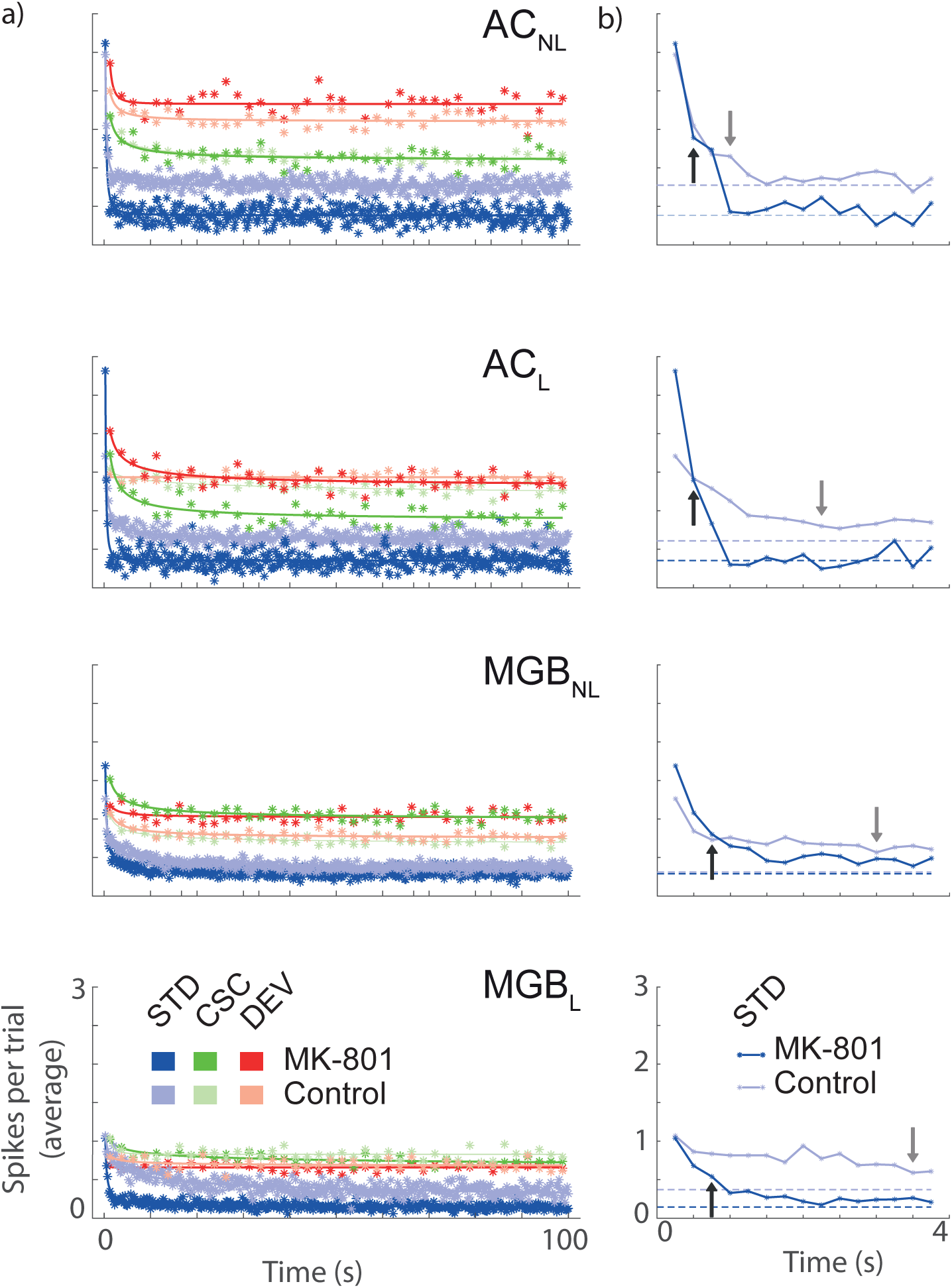
Time course for dynamical thalamocortical adaptation. **a)** Averaged time course for the stimulus played in relation to the time elapsed from the beginning of the sequence. **b)** The first fifteen standard stimuli showing the three parameters of the power low fitted: *a* initial average response; *b* adaptation velocity; and *c* the steady-state value (dotted lines) for each group. Arrows represent the 50% of the initial responses demonstrating faster adaptation in the MK801 group and the break down in the dynamical hierarchy of adaptation.

These data reveal that MK-801 alters the timing across the hierarchical organization of the auditory system, resulting in the lemniscal thalamus having almost the same adaptation velocity as the non-lemniscal cortex (arrows in Figure 4b). Furthermore, MK-801 reduces (almost by half) the steady-state plateau in the AC (dotted lines in Figure 4b; *c* values for control group: MGB_L_=0.0776, MGB_NL_=0.2908, AC_L_=0.6084 and AC_NL_=0.7740; and for MK-801 group: MGB_L_=0.1428, MGB_NL_=0.2884, AC_L_=0.3523 and AC_NL_=0.3834).

All these results together support the idea that MK-801 produces a differential effect on adaptation and deviance detection along the thalamocortical axis, providing new evidence of a change in the firing pattern and temporal responses at single neuron level.

### Delayed and broader larger-scaled LFP responses

Next, we wanted to check if the single unit responses correlated with larger-scale measurements of neuronal activity. The analysis of local field potentials (LFP) revealed that MK-801 produced significant changes in MGB_NL_ and AC (both in the lemniscal and non-lemniscal portions) for the deviant, standard and cascade LFPs (Figure 5a-c), such that they exhibited broader and longer responses for DEV-LFP and CAS-LFP in the auditory cortex, while the waveforms of these LFPs were shifted in latency for the MGB_NL_ due to a progressive delay of N1, P1 and N2 (note that this terminology refers to the first negative peak, first positive peak and second negative peak), showing delayed peaks of 8, 14 and 57ms for DEV-LFP and 6, 26 and 45ms delay for CAS-LFP in N1, P1 and N2, respectively (DEV-LFP: N1 peak for MK-801= −6.6μV at 20ms and control= −1.5μV at 12ms; P1 peak for MK-801=6.9μV at 41ms and control=6.8μV at 28ms; finally, N2 peak for MK-801= −5.4μV at 102ms and control= −10.2μV at 45ms. CAS-LFP: N1 peak for MK-801= −6.5μV at 18ms and control= −1.2μV at 12ms; P1 peak for MK-801=5.1μV at 53ms and control= 10.6μV at 27ms; finally, N2 peak for MK-801= −5.0μV at 91ms and control= −10.3μV at 45ms).

**Figure 5.**
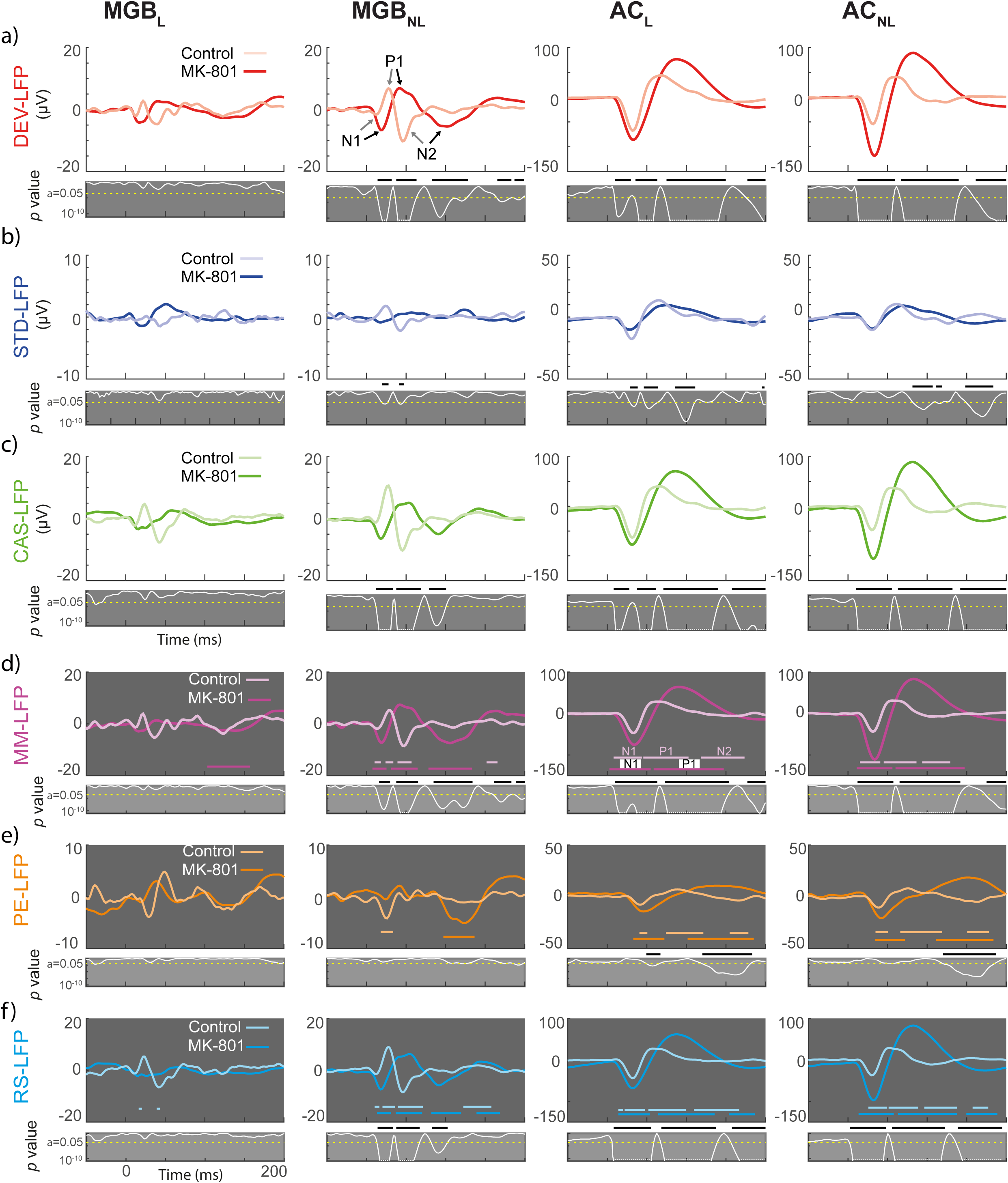
Local Field Potentials for each condition and their differences. **a-c)** Population grand-averaged LFP for each condition recorded (CAS, DEV and STD) within each group (controls and MK801). Grey panels under the main LFP representations shows the instantaneous *p* value (white trace) of corresponding stimulus condition LFP (critical threshold set at 0.05 represented as a horizontal dotted yellow line). The thick black horizontal bars in figure 5a-c highlights the time interval for which the LFP comparison between the control and MK801 groups is significant. **d-f)** Population grand-averaged LFP for and neuronal Mismatch (MM-LFP=LFP_DEV_-LFP_STD_), Prediction Error (PE-LFP=LFP_DEV_-LFP_CAS_), and Repetition Suppression (RS-LFP=LFP_STD_-LFP_CAS_) respectively Colored horizontal lines denote significative deflections (*t*-test, FDR corrected). Grey panels show the instantaneous *p* value (white trace) of corresponding stimulus condition LFP (critical threshold set at 0.05 represented as a horizontal dotted yellow line) and black horizontal lines the time interval in which MK801 and control are statistically different.

**Figure 6.**
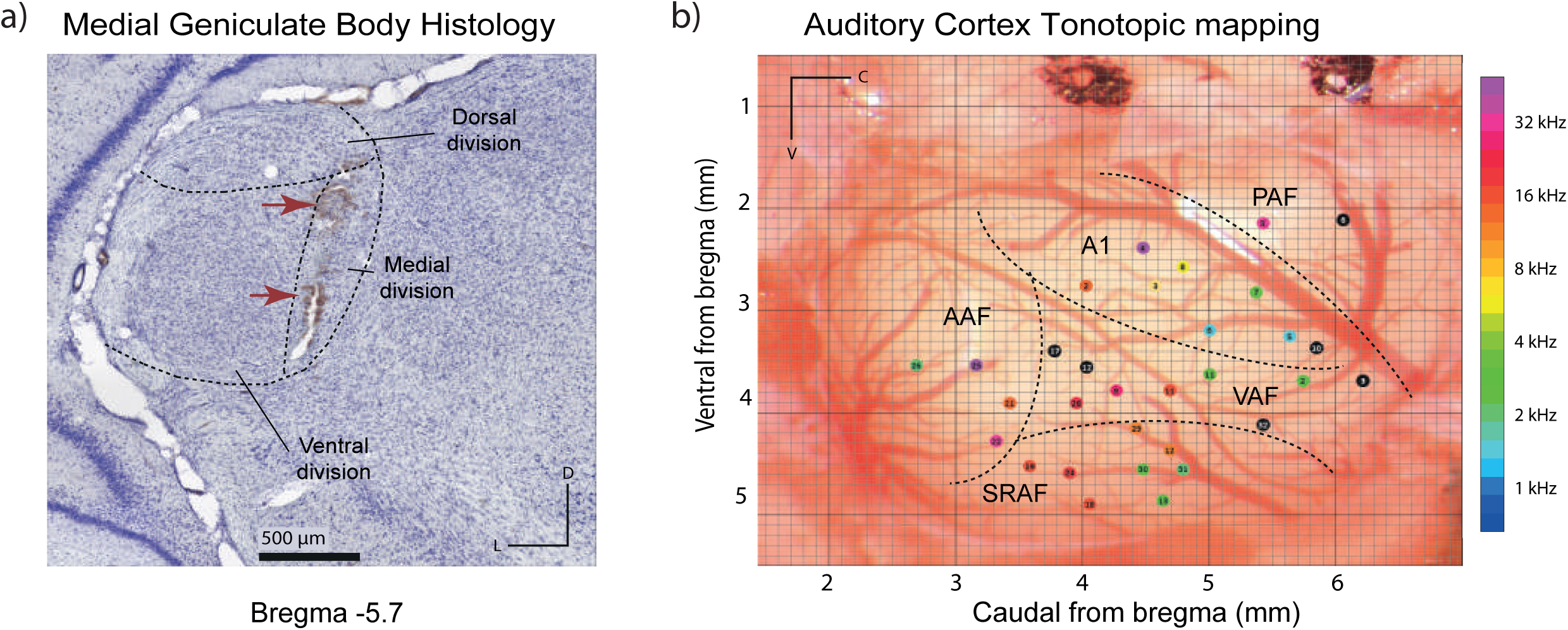
Anatomical recordings location. **a)** Photomicrography sample of a MGB Nissl-stained slice (10x), red arrows point the two electrolytic lesions. **b)** Example of localization all recordings made in the AC of one rat, each colored dot represent the characteristic frequency of each performed tract. A1: Primary Auditory Field; AAF: Anterior Auditory Field; VAF: Ventral Auditory Field; PAF: Posterior Auditory Field and SRAF: Suprarhinal Auditory Field.

Similarly, we also sought significant LFP signals for each computed index (Figure 5d-f). The horizontal colored lines highlight the time at which significant deflections occur to each index-LFP for control and MK-801 groups independently (light and bright horizontal lines, respectively). Additionally, we compared these LFP indices between groups. The analysis of the MM-LFP shows that MK-801 elicited stronger and broader deflections within all regions (horizontal bright purple lines; Figure 5d) and abolished the late negative component (N2) in the AC (MGB_L_: N2 = 114-157ms; MGB_NL_: N1 = 12-21ms, P1 = 32-63ms and N2 = 75-135ms; AC_L_: N1 = 10-57ms and P1 = 60-147ms; AC_NL_ N1 = 20-53 and P1 = 60-144ms). Our data also demonstrate that MK-801 produced a higher MM-LFP for virtually the whole LFP response within MGB_NL_ and both AC, while no differences occurred in MGB_L_.

Similar to the spike population analysis, and considering that the PE-LFP and RS-LFP both contribute to the MM-LFP, we also wanted to understand how MK-801 shapes the LFP for prediction error and repetition suppression. In response to MK-801, the PE-LFP waveform was reduced at the early component of the MGB_NL_, while it was increased and delayed for the AC (orange horizontal lines in Figure 5e). Moreover, MK-801 also abolished the N2 deflection (MGB_NL_: N1 = 99-146ms; AC_L_: N1 = 30-65ms and P1 = 87-180ms; AC_NL_ N1 = 30-67 and P1 = 106-180ms). When PE-LFP was compared between groups, we only found differences in AC, mainly at the early (50-70ms) and late components (120-180ms). In other words, the lemniscal thalamus does not exhibit deviance detection, neither at the single neuron level nor at large-scale responses. Hence PE-LFP confirm single unit population data, where MK-801 produced greater levels of deviance detection in the auditory cortex (figure 2e).

Finally, MK-801 had similar effects on RS-LFP to those described above for MM-LFP and PE-LFP, eliciting broader and larger waveforms for MGB_NL_ and AC (Figure 5f; MGB_NL_: N1 = 10-28ms, P1 = 34-63ms and N2 = 73-108ms; AC_L_: N1 = 10-55ms, P1 = 67-140ms and N2 = 148-180ms; AC_NL_: N1 = 10-51, P1 = 55-132ms and N2 = 141-180ms). When differences between groups are considered, the non-lemniscal thalamus exhibited a shift in the waveform between 15-100ms, while for the cortex, responses over virtually the whole temporal window were increased by MK-801.

## DISCUSSION

In this study, we demonstrate that the neuronal index derived from single cell recordings of mismatch is profoundly affected along the auditory thalamocortical system in rats treated acutely with a low dose of the NMDA-R antagonist, MK-801. Importantly, we also reveal that the two elements that make up the index of mismatch negativity, i.e., repetition suppression and prediction error, are differentially affected by MK-801 in single neurons at auditory thalamus and cortex. MK-801 increases repetition suppression in thalamus and prediction error in cortex. The increase in repetition suppression is more prominent in lemniscal areas of the thalamus, while the increase in prediction error is more evident in the non-lemniscal areas of cortex. Furthermore, our results demonstrate that MK-801 alters the dynamics of neuronal adaptation along the thalamocortical axis, becoming faster and stronger especially at thalamic level. The pattern of results from single unit data were confirmed by recordings of large-scale responses, LFPs, as the latter exhibit delayed and broader deflections. In summary, our work demonstrates that the MK-801 increase of the neuronal mismatch in the auditory cortex 60ms after stimulus onset is due to the combined effect of an increment in the sustained responses to deviant tones and a decrement to standard tones. It should be noted that, in contrast to most previous studies using recording procedures to study neuronal population activity in rodents such as LFPs or EEG via skull screws, we have recorded single-unit activity, a technique that has a much higher resolution level at the single cell level for revealing patterns of activity underpinning mismatch responses.

It is well established that NMDA-R plays a fundamental role in neuronal plasticity, controlling long-term potentiation and depression (Blanke & VanDongen, 2009). Further, it is generally accepted that human MMN is reduced after NMDA-R antagonist treatments because NMDA-R antagonist blocks synaptic plasticity, precluding the formation of a memory trace for the standard tones (Todd *et al.*, 2013). As we have seen in our results, MK-801 reduces responses to standard tones thus increasing repetition suppression.

Although this finding supports the hypothesis that NMDA-R antagonists alter sensory-memory formation (Auksztulewicz & Friston, 2016), the findings that low dose (0.1 mg/kg) MK-801 treatment produces a significant increment in the response to the deviant tones, in prediction error and hence, an increment in the neuronal mismatch, are in the opposite direction to expected. It is clear that the role of NMDA-R in the generation of MMN is considerably more complex than thought (Harms *et al.*, 2018). There have been suggestions in the literature of precedents for our observations. Even considering that MK-801 has 160 times the affinity of ketamine to NMDA-R, necessitating higher ketamine doses for similar drug effect (Schuelert *et al.*, 2018), our results conform with those that report an increment in amplitude and latencies to deviant responses after the acute ketamine treatment in rats (Ahnaou *et al.*, 2017) and with a sub-anaesthetic dose of ketamine in healthy humans producing larger N100 to deviant tones but not MMN (Oranje *et al.*, 2000). Interestingly, a dose response study of the MK-801 effects on MMN-like responses in male rats showed that while a high dose (0.5mg/kg) reduced late deviance detection (around 55ms), a medium dose (0.3mg/kg) significantly enhanced early deviance detection effects (at about 13 ms) and some evidence of enhanced late effects although not significantly (Harms *et al.*, 2018). We used a single dose of 0.1mg/kg in female rats, as it has been demonstrated that females are more sensitive to MK-801 than males (Andine *et al.*, 1999) and that this dose is enough to induce behavioral effects (Meehan *et al.*, 2017). Importantly, memantine, a low affinity uncompetitive agonist of NMDA-R, has been shown to (i) increase the duration of rodent MMN-like responses (Tikhonravov *et al.*, 2010), (ii) increase MMN amplitude in healthy individuals (Korostenskaja *et al.*, 2007), and (iii) in persons with schizophrenia (Swerdlow *et al.*, 2016).

The memantine results suggest an interpretation of our findings in terms of the mechanisms underpinning synaptic plasticity (Slutsky *et al.*, 2004). Partial blockade of NMDA-R channels (such as mediated by memantine, or low dose MK-801) is also likely to reduce background calcium flux resulting in homeostatic upregulation of NR2B-containing NMDA-Rs leading in turn to the conversion of synapses to a plastic state. That is, while these drugs reduce calcium influx during uncorrelated activity, there is increased calcium influx during correlated activity (produced by physiological stimuli), increased signal to noise, facilitated transmission and increased plasticity (Slutsky *et al.*, 2010).

Other characteristics of the neuronal mechanisms and microcircuitry involving the glutamate NMDA-R system are relevant to the effects we have observed on the neuronal mismatch after the MK-801 treatment. NMDA-R are located, not only at postsynaptic and presynaptic sites in excitatory neurons, but they are also found at GABAergic inhibitory interneurons in neocortex (DeBiasi *et al.*, 1996). MK-801 has demonstrated a preferential regulation of the firing rate of cortical GABA interneurons, increasing the firing rate of the majority of pyramidal neurons (Homayoun & Moghaddam, 2007) and therefore producing an imbalance in the excitatory/inhibitory networks in the cortices (Bygrave *et al.*, 2016; Okada *et al.*, 2019). It is well known that cortical GABAergic interneurons differentially amplify stimulus-specific adaptation (a similar phenomenon to iMM) in excitatory pyramidal neurons in auditory cortex (Chen *et al.*, 2015). Moreover, a model of a mutually coupled excitatory/inhibitory network can explain distinct mechanisms that allow cortical inhibitory neurons to enhance the brain’s sensitivity to deviant or unexpected sounds (Natan *et al.*, 2015). Further MK-801 would alter the tonic inhibitory control of NMDA-R in cortical areas leading to the activation of pyramidal neurons by subsequent deviant tones.

The increased repetition suppression we observed in the medial geniculate body could also be by altered excitatory/inhibitory balance. Although the rat MGB lacks GABAergic neurons, it receives GABAergic input from the thalamic reticular nucleus (TRN) and the inferior colliculus (Malmierca, 2003). The latter is a source of bottom-up inhibitory influences while the TRN provides the MGB with an indirect and inhibitory feedback activation from AC (Bartlett, 2013). Cortical stimulation hyperpolarizes TRN neurons and increases their inhibitory output to the MGB (Crabtree *et al.*, 2013) and furthermore, TRN has been demonstrated to profoundly influence SSA in the MGB (Yu *et al.*, 2009). Changes in the thalamocortical neuronal firing pattern of thalamic neurons into bursts have been suggested to provide an alerting signal to the cortex to enhance stimulus detection (Hu & Agmon, 2016). Overall our results match the general concept that when the system is adapted, it is more sensitive to detect changes in the environment (Musall *et al.*, 2014), where a stronger thalamic repetition suppression (or inhibition) support the increase in the prediction error signals (excitatory) at cortical level, or vice versa. It would be interesting to test whether thalamic repetition suppression is correlated with cortical prediction error signals, but this question awaits future experiments.

Our study is important because it has revealed the involvement of two basic mechanisms, i.e., repetition suppression and prediction error; and two different pathways, lemniscal and non-lemniscal, underlying the neuronal mismatch in the thalamocortical hierarchy. Predictive coding theory proposes that the brain constantly tries to minimize the discrepancy between actual sensory input and internal representations of the environment (Friston, 2005). What is new in our data is the critical importance of the hierarchical organization of the auditory system in sharing the ‘responsibility’ for generating the representation and detecting the discrepancy, largely attributable to thalamic and cortical processes. However, our data provide evidence that the NMDA-synaptic plasticity and MMN relationship is not as simple as previously surmised from human studies. Admittedly, we have only tackled the functional role of the NMDA-R under a particular experimental manipulation and we cannot exclude the possibility that larger doses of MK-801 would have generated different results. It is also well known that other neuromodulatory systems such as the dopaminergic, cholinergic and/or cannabinoid systems may be altered and interact with the NMDA-Rs in normal brain function (Ayala *et al.*, 2016; Valdés-Baizabal *et al.*, 2017) as well as in schizophrenia (Howes & Kaar, 2018; Lucatch *et al.*, 2018; Parr & Friston, 2018; Okada *et al.*, 2019). What are the implications of our findings for schizophrenia? Reduced MMN is associated with poor global functioning (Light & Braff, 2005) and cognitive deficits (Baldeweg *et al.*, 2004) in schizophrenia. Hence, if a safe drug were available that targeted the relevant NMDA-R subunit, and facilitated neuroplasticity as indexed by increased MMN, it offers opportunities for interventions to remediate cognitive deficits that are a core feature of schizophrenia (Green *et al.*, 2000). Memantine which has been shown to increase MMN amplitude in healthy individuals and in schizophrenia has been used as an adjunctive therapy in schizophrenia for some time to improve cognition in particular. While effects of adjunctive therapy are small, recent meta-analysis suggests that there are improvements in global measures of cognition, but improvements in more sensitive composite cognitive test scores have not been observed (Kishi *et al.*, 2018). To date, there have been no attempts to utilize MMN enhancement to memantine as an index of increased neuroplasticity that could be exploited in remediation studies. Interestingly, both the moderate affinity antagonist, memantine, and high affinity antagonist, MK-801, bind to the NR2B subunit of the NMDA-R at very similar binding locations (Song *et al.*, 2018) but only memantine has been approved for use in humans given evidence of neurotoxic effects of MK-801 in humans (Olney *et al.*, 1989). One avenue of future research is the development of safe compounds for human use that target similar binding locations to memantine and MK-801.

### Data availability

The data that support the findings of this study are available from the corresponding author on reasonable request.

## Acknowledgements

We thank Drs. Javier Nieto-Diego and David Perez-Gonzalez for their technical support and constructive criticisms; and Drs. Nashat Abumaria and Juanita Todd for advice and constructive comments on the discussion. We also thank Ms. Marianny Janiree Pernía Rosales for helping with the histological photomicrography and Mr. Antonio Rivas Cornejo for the technical support.

